# A Model For Hierarchical Memory Storage in Piriform Cortex

**DOI:** 10.1101/2022.10.27.514001

**Authors:** Achint Kumar

## Abstract

How the brain stores and retrieves memories is an important unsolved problem in neuroscience. It is commonly believed that memories are represented in the brain by distinct patterns of neural activity. Attractor dynamics has been proposed as one of the theoretical frameworks for learning and memory in neural networks. However, most of the prior theoretical work typically assumes that the neural network consists of fully-connected, binary neurons and that neuronal representations of memories are uncorrelated. In this paper, we propose a model consisting of continuously varying, rate-based, sparse neural network with a local learning rule which stores correlated patterns organized into multiple uncorrelated classes. We perform analytical calculations to compute maximum storage capacity and basin of attraction. It is found that increasing pattern correlation decreases storage capacity, and increasing the memory load decreases basin of attraction. We also study rate-based and spiking based neural network with separate excitatory and inhibitory populations and tight EI balance. Recent experimental work indicates that piriform cortex exhibits attractor dynamics and possibly stores hierarchically correlated patterns. So, we consider this work as a model for olfactory memory storage.

## 1 Introduction

Understanding memory formation and learning in the brain is one of the central goals of neurobiology. Investigating olfactory processing is particularly well suited to begin to achieve this goal for a couple of reasons. First, olfactory cortex has a simpler 3 layered architecture as opposed to the 6 layered architecture of neocortex. Second, it is much shallower than other sensory modalities. The receptor neurons and cortical neurons are separated by just two synapses. It is hoped that the techniques developed and computational principles discovered in the process of understanding this simple system could be modified and generalized to understand other sensory modalities and cognitive processes of the brain.

In this paper, we investigate memory formation in a particular area of olfactory cortex known as the piriform cortex(PCx). Multiple sources of evidence points to piriform cortex as being the site for odor memory storage[1]. Experimentally, lesions of PCx or afferent input to PCx results in impairment of odor discrimination ability [2, 3] and theoretically, the diffuse input and recurrent connectivity in PCx are highly reminiscent of abstract memory models [4].

An animal foraging in an environment should be able to recognize a previously encountered food item even if the relative concentration of different odorant molecules making it up are not the same as when they were first encountered. This stability of representation of an odor object (like apple) to small variations in concentration of constituent odorants is called pattern completion. An animal must also be able to discriminate between two food items with highly overlapping constituent odorants (like fresh orange and rotten orange). This separation of closely related olfactory stimulus is called pattern separation. Attractor dynamics is a framework for realizing pattern separation and completion in neural networks. In this paradigm, the neural network has specific stable states which are stable fixed points (called attractors) that attract nearby states.

Recent work suggests that piriform cortex accomplishes pattern separation and completion through attractor dynamics [5, 6]. The most popular model of this activity is known as the Hopfield model which uses a Hebbian learning rule to update synaptic weights [7]. Hebbian learning corresponds to a local learning rule in which synchronous activation of two neurons increases the synaptic strength between them and asynchronous activation decreases it. The use of this learning rule to store different patterns shapes the synaptic connectivity of the network, leading to the formation of an energy landscape with valleys representing corresponding attractors. The network evolves in time to descend the energy landscape to the nearest valley. Such networks are characterized by strong recurrent synapses (compared to feedforward and feedback connections) and demonstrate persistent activity even after the stimulus is ceased. Neurophysiologically, these patterns are assumed to correspond to different memories stored in the brain. In addition to potentially explaining the functionality of piriform cortex, attractor dynamics have been used to explain previous experimental work regarding persistent activity of certain neurons in monkeys [8, 9] and mice [10] during working memory tasks.

For simplicity, in the Hopfield model the stored patterns are assumed to be uncorrelated. Crucially, the model fails in storing correlated patterns as the network ends up retrieving a combination of stored patterns. This shortcoming is especially important for memory storage in the piriform cortex, as it has been recently found that memories stored in PCx are retrieved by neurons firing in correlated patterns for different odors [11]. Likewise, it is highly possible that PCx stores disparate olfactory memories as correlated patterns, and current models are not able to reproduce this type of memory storage in an efficient and biophysically realizable manner.

In this paper, we consider a theoretical toy model for storing hierarchically organized memories in a rate-based neural network evolving via a local learning rule. Our model categorizes the different memory patterns into various disjoint classes(see Fig 1a). Each class is represented by a prototype which is uniformly correlated with the exemplar patterns in that class. However, the members from different classes are not correlated which leads to a block diagonal correlation matrix structure(see Fig 1b). The original Hopfield prescription[7] allows us to store only weakly correlated patterns. Our proposal, which is a generalized form of a previously proposed model allows us to store strongly correlated patterns exhibiting block diagonal correlation structure[12]. This paper is organized as follows. First, we present a learning rule which allows us to store hierarchically organized patterns. Next, we show how neuronal transfer function parameters affects decorrelation. Then, we discuss the results of mean field analysis and show that increasing correlations decreases maximum storage capacity. Then we look at the effect of memory load on basin of attraction which is experimentally testable by silencing certain glomeruli in olfactory bulb and measuring changes in neural responses in piriform cortex.

**Figure 1:**
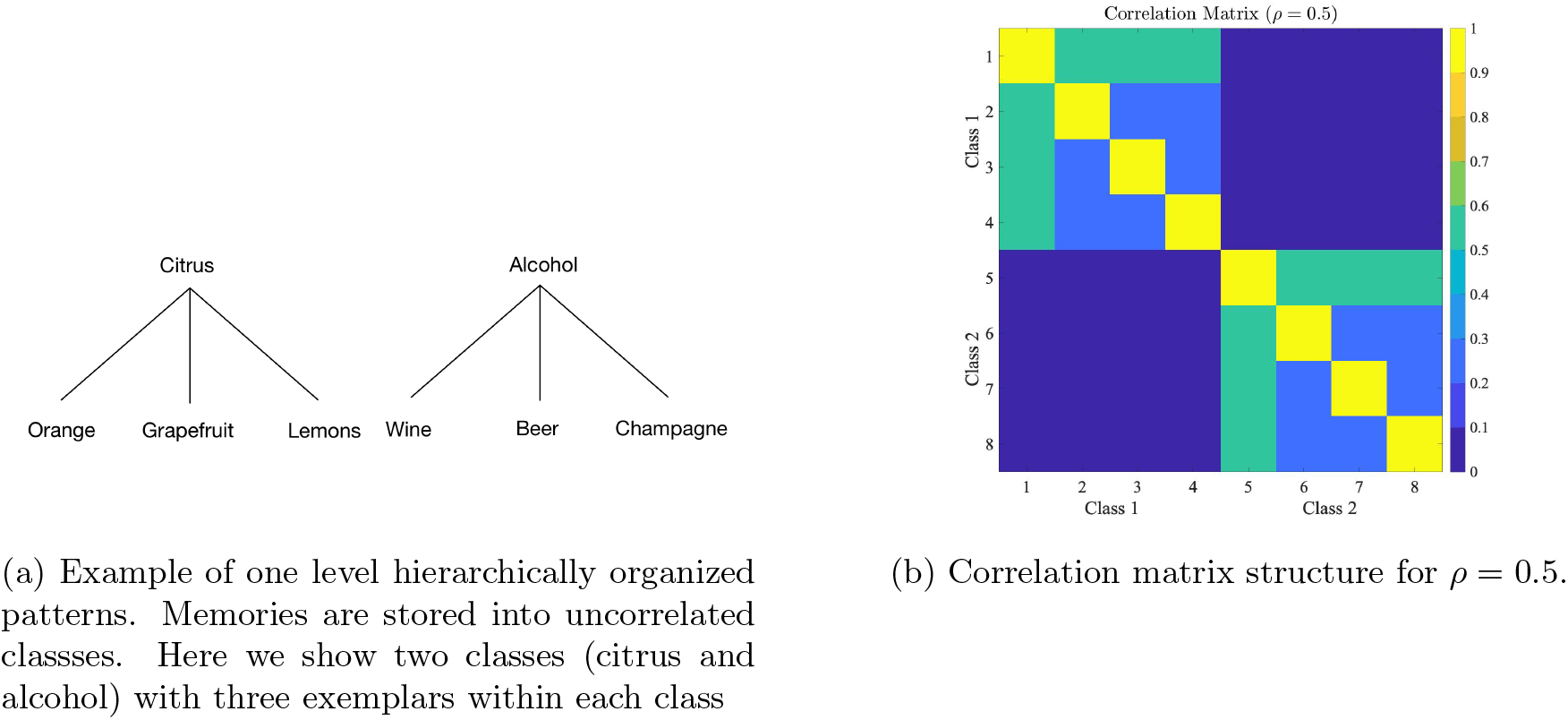
Hierarchically organized patterns

## 2 Problem Setting

Piriform cortex consists of multiple excitatory and inhibitory neuronal populations connected to each other in ways that have not been completely mapped out. These neurons receive input from olfactory bulb, other parts of the olfactory cortex and various cortical areas of the brain. We start by assuming that all the neurons in our model of piriform cortex receives input from the olfactory bulb only, and that the recurrent neural connectivity is spatially homogeneous. Let us further ignore the distinction between excitatory and inhibitory neurons at this point. For analytical tractability, we begin by considering rate-based network of N neurons evolving according to standard dynamical equations[13]

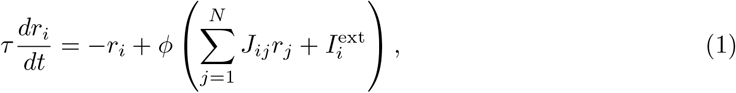

where *τ* is the time constant of rate dynamics, *r*_*i*_ *>* 0 (*i* = 1, …, *N*) is the firing rate of *i*^th^ neuron, *J*_*ij*_ is the weight matrix, 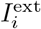 is the external (olfactory bulb) input to *i*^th^ neuron, and *ϕ* is a sigmoidal neuronal transfer function,

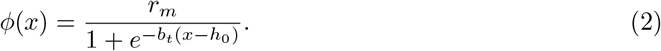

This form of transfer function has been earlier shown to fit data[14]. The parameters *r*_*m*_, *b*_*t*_ and *h*_0_ were carefully chosen so that the network shows attractor dynamics(see Table 1 of SI).

## 3 Results

### Decorrelation using functional non-linearity

We begin by assuming that the olfactory stimulus space is tiled by *P*_*C*_ classes with *P*_*I*_ exemplars within each class. The class prototype is represented by 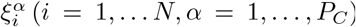 and exemplars of *α* class are represented by 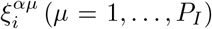. We assume that the different class prototypes and exemplars are uncorrelated but there exists uniform correlation between exemplars and their class prototype given by *ρ*. The correlation structure between class prototypes and exemplars is given by 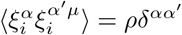. Since, the output of neuron i is 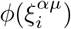, the usual weight matrix by Hebbian rule should be of the form: 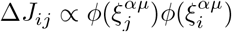. But it turns out a better fit to neural data is provided by taking into the affect on synaptic strengths to various inputs. So, a better alternative turns out to be 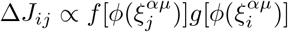 where both f and g have a functional form 1*/*2(2*q* − 1 + tanh[*β*(*x* − *x*_0_)]) whose the parameters *q, β* and *x*_0_ are carefully chosen so that they closely fit experimental data[15, 14].

An additional affect of these nonlinearities is that it the correlation between the exemplars and prototype decreases. We denote the new correlations by *b* so we have 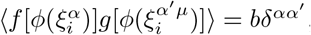,. The relationship between b and *ρ* can be analytically calculated (see SI) and it turns out that increasing threshold and sparsity increases decorrelation (see Fig 2).

**Figure 2:**
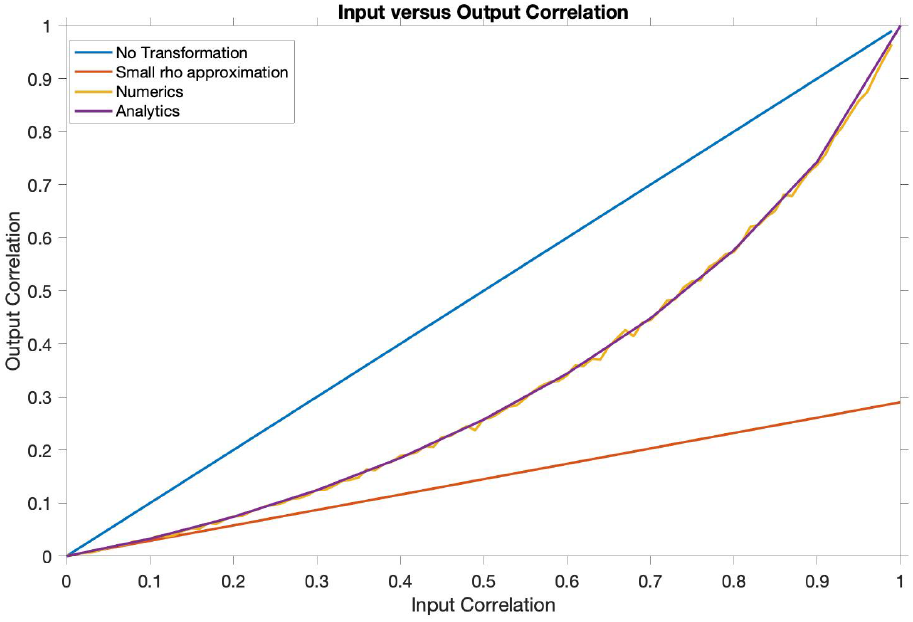
Input versus Output correlation brought about by (*f* [(*ϕ* (*ξ*))] transformation. We show the results of small rho approximation(SI), analytical formula and the results of simulations. The blue line indicates no transformation

### 3.1 Learning Rule

Our synaptic weight matrix is inspired from previous works [12] that have proposed a learning rule prescription to store hierarchically organized patterns in fully-connected network of binary neurons. Here, we generalize that prescription and make our network biologically more realistic by considering a sparse network consisting of rate units, and storing Gaussian patterns. Since most neurons in the piriform cortex receives hundreds of input from other neurons, then using Central Limit Theorem, it is not unreasonable to suppose that the net input to any neuron is sampled from a Gaussian distribution.

The weight matrix is given by

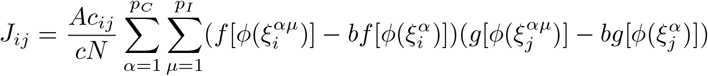

where A controls the strength of recurrent connectivity, *c*_*ij*_ controls the sparsity of the network and is drawn from i.i.d. Bernoulli random variable (*c*_*ij*_ = 1 with probability c and 0 otherwise).

### 3.2 Learning Hierarchically Correlated patterns

With the generalized weight matrix, we simulated three different networks. The first network assumed a homogeneous population of neurons. We were able to find persistent activity in this network by carefully choosing parameters (see Table 2 of SI). The next network we simulated assumed a separated EI rate network (see middle of Figure 3) and separated EI spiking network (see right of Figure 3).

**Figure 3:**
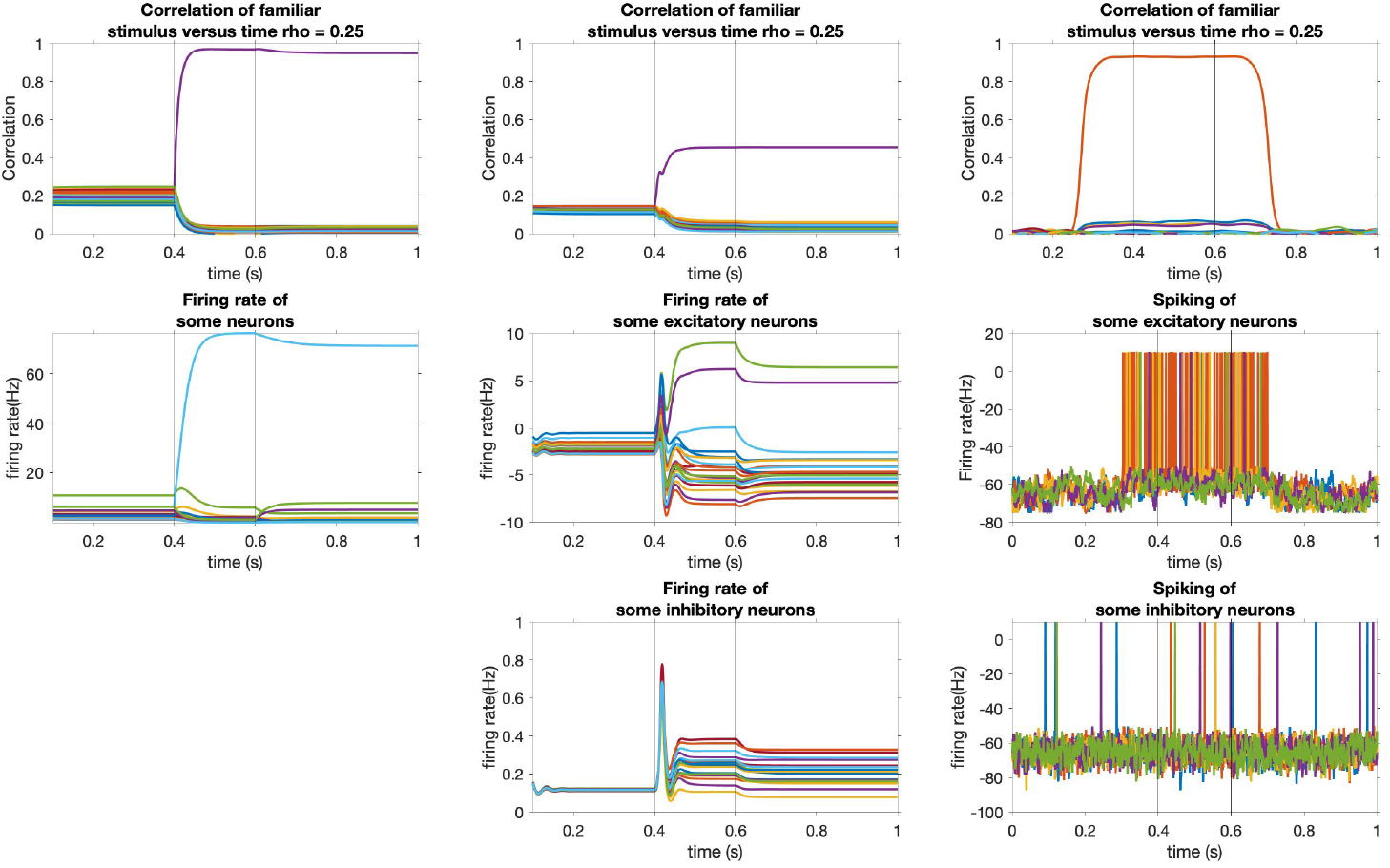
Persistent activity is a signature of attractor dynamics. The vertical lines represent the onset and offset of the presented stimulus.**Top** : Correlation with familiar stimulus versus time for rate, rate EI, and spiking network. **Middle** : Middle Firing rate of some random neurons(left) and excitatory neurons(middle), spiking of some excitatory neurons(right). **Bottom** : Firing rate of some inhibitory neurons(left) and spiking of some inhibitory neurons(right)

### 3.3 Storage Capacity

Next, we performed mean field analysis to get an exact expression for the overlap *m* of the response of the network to the presented pattern as a function of correlation between the memory classes. The result is shown in Figure 4. In Figure 4a, we see that the steady state overlap of network with presented memory decreases faster if the correlations are larger. We also notice that for high correlations, *ρ* = 0.6, 0.7 in the figure, there is no retrieval. In Figure 4b, we show how the memory capacity decreases with increase in correlation. The model shows a second order phase transition at around *ρ* = 0.54.

**Figure 4:**
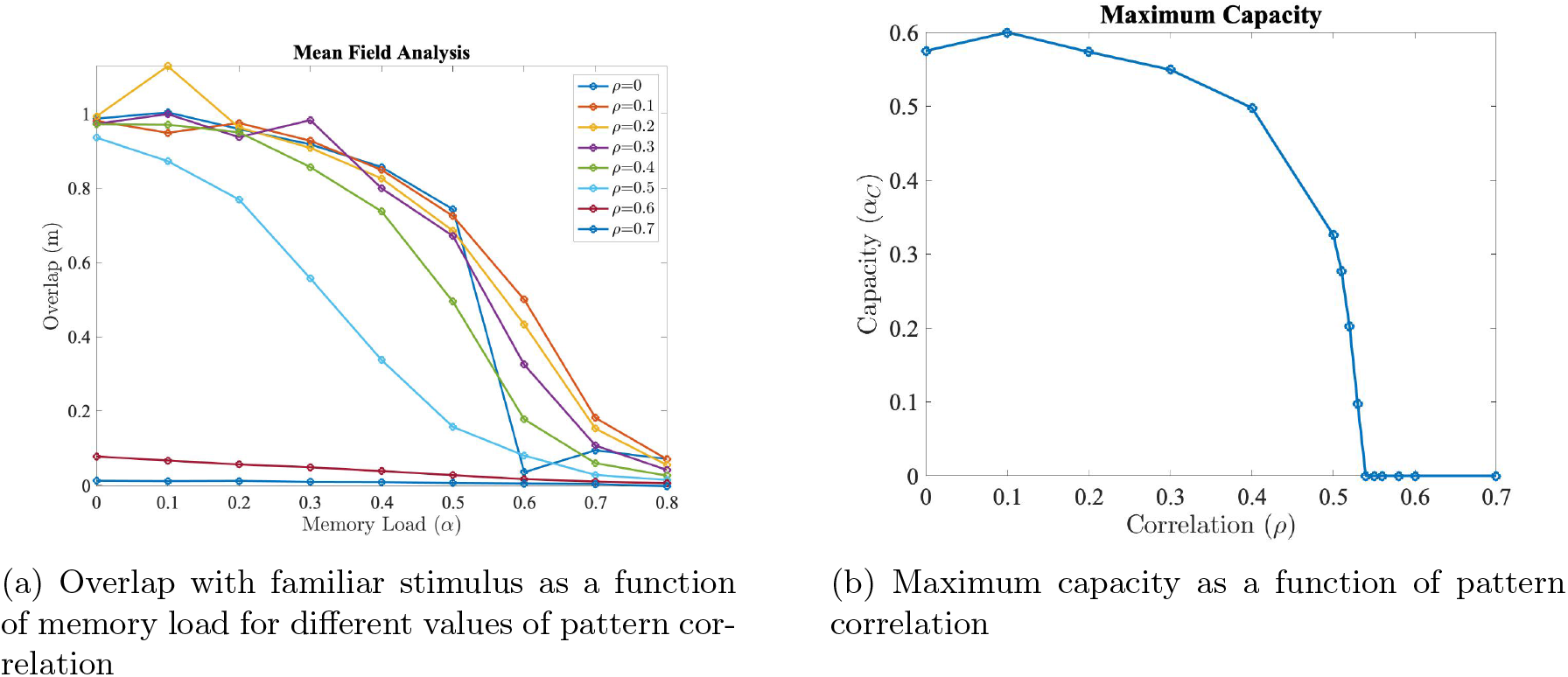
Mean Field Analysis of the learning model

### 3.4 Basin of Attraction

The basin of attraction of stored memories is measured by varying the amount of initial overlap with a stored memory and measuring the time evolution. Suppose we would like to retrieve 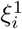, then we start with an initial state of the network, 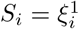 for *i* = 1, 2, 3 …, *mN*, while for *i > mN, S*_*i*_ is chosen at random from a gaussian distribution. For large values of m, network doesn’t flow to *ξ*_1_ but as m decreases a substantial number of iterations leads to a flow towards *ξ*_1_. This process is repeated for other initial states corresponding to different stored patterns. The mean of the overlap of the final network state with the different stored patterns whose corrupted version was presented to the network is calculated. The largest value of m corresponding to the final overlap with the stored pattern*>* 0.5 gives us the radius of attraction. We repeat the procedure for different values of memory load. In Figure 5 (top), we see the network state overlap with presented memory as a function of correlation. We see expect decrease in overlap as the memory load increases. In Figure 5 (bottom), we see that the radius of attraction as a function of memory load. There is a sudden drop in radius of attraction between *ρ* = 0.4 and *ρ* = 0.5 indicating there is a first order phase transition.

**Figure 5:**
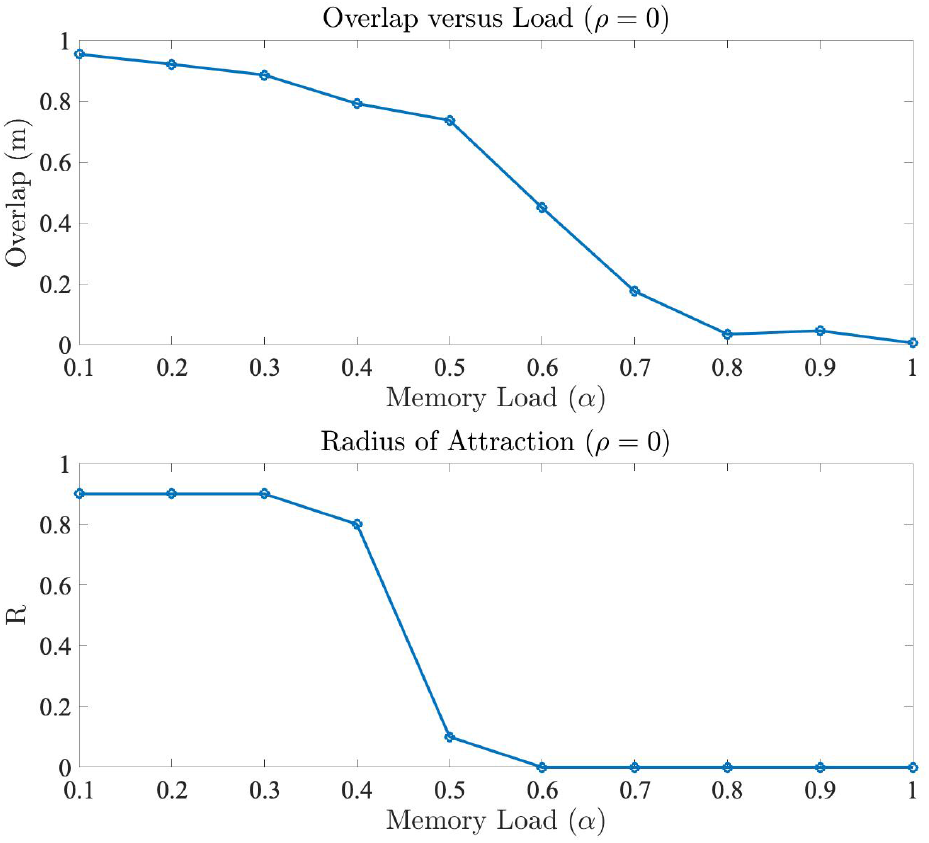
**Top** : Overlap as a function of memory load for uncorrelated patterns. **Bottom** : Basin of attraction for different values of memory load

## 4 Discussion

Multiple experimental evidence supports the hypothesis that PCx shows attractor dynamics with correlated representation for different odors. In this paper, we described a theoretical model that prescribes a possible way to store correlated associative memories in a neural network. We showed the existence of second order phase transition in maximum capacity of the network as a function of memory correlations using mean field analysis. We also showed that the basin of attraction undergoes first order phase transition.

While our model allows us to store correlated patterns using a local learning rule, it is unclear exactly how the different terms in our learning rule can implemented in a biological system. In addition, piriform cortex has a spatially non-homogeneous connectivity, multiple populations of excitatory and inhibitory neurons, and receives time-varying input. Our proposed model accounts for none of these features. Despite these limitations, we believe that our model serves as a starting point towards a more detailed understanding of olfactory memory formation.

On the experimental side, we need to understand exactly how the neural interactions lead to storage of odor representations in the PCx. One way to investigate this would be to perturb the activity in the olfactory bulb using optogenetics and record from PCx and see if the odor representation remains invariant under these perturbations. A prediction of attractor dynamics is that small amount of perturbation would lead to stable representation in PCx but once the perturbation exceeds a critical limit, the network would leave the basin of attraction and evolve into a completely different state. This experiment would allow us to test the existence of first phase transition that is predicted by our model. The role of PCx recurrent connectivity in memory formation can be tested by developing novel toxins which can selectively inhibit those connections and studying the modified response to odorants.

## 5 Materials and Methods

Details of the numerical and analytical calculations are in the Supplementary Information, including the mean-field theory, construction of excitatory-inhibitory rate network, and details of the LIF spiking network. The simulations were performed using MATLAB and Mathematica. All the code is available on GitHub, https://github.com/achintzeus1994/CorrelatedPatterns. We thank Nicolas Brunel and Kevin Franks for helpful discussions in the initial phase of this project.

## Supplementary Information

### 6 Model

A sparse rate-based neural network consisting of N neurons is considered. The dynamics of the neurons is given by the following standard system of differential equations:

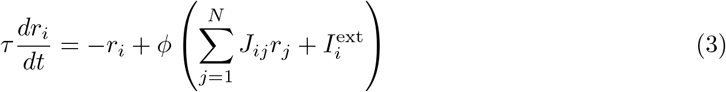

Here *r* ≥0, is the firing rate of the neurons and *τ* is their time constant which is assumed to be the same for all neurons. The external input to neuron *i* is 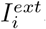. Here *ϕ* is the single neuron transfer function and has the following functional form:

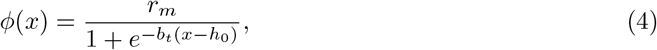

The values of the parameters *r*_*m*_, *b*_*t*_ and *h*_0_ are carefully chosen to give rise to attractor dynamics. Their(and other parameters) precise values are listed in Table 1.

The neural network is designed to store hierarchically correlated patterns. Previously, two different prescriptions have been proposed which allows us to store such patterns in binary, fully-connected neural network. [16, 17]. We generalize those prescriptions to store continuous patterns in a sparse network with rate-based units. We assume that there are *p*_*C*_ classes represented by prototype patterns 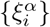 where *α* = 1, 2…*p*_*C*_. Each class has *p*_*I*_ exemplar patterns denoted by 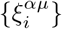 where 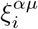 can be thought as *µ*^*th*^ pattern in class *α* input to neuron i. The two models investigated are:

- Gutfreund Model: This model was specifically designed to store hierarchically organized patterns. The generalized weight matrix prescription we use in this paper is given by:

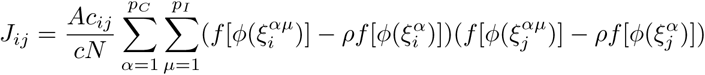

Here A is the learning rate, *c*_*ij*_ are i.i.d. Bernoulli random variables(*c*_*ij*_ = 1 with probability c and 0 otherwise) and *ρ* is the correlation between the exemplar and prototype of that class. The functions f(.) and g(.) are taken to be of the form:

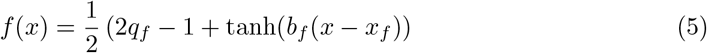

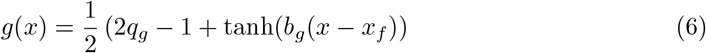

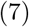

The parameters of f(.) have previously been constrained by data from ITC cortex. The parameter values we use are listed in Table 1.
- Pseudo-Inverse Model: Unlike the previous model, the patterns don’t need to be hierarchically organized in this framework. So, this model provides a generalized framework for storing correlated patterns. However, since we are interested in hierarchically stored patterns only, we limit our analysis to that particular case. The key feature of pseudo-inverse model is that the stored patterns are the fixed points of the network. So mathematically we would like, 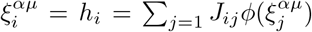, where *h*_*i*_ is the input to the *i*^*th*^ neuron. The following weight matrix prescription satisfies this constraint:

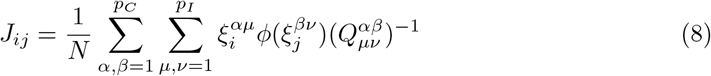

Here Q is the overlap matrix which is given by:

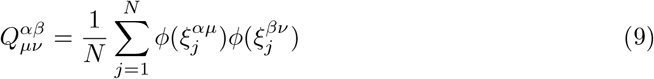

The advantage of Gutfruend model is that it is local while, in general the pseudo-inverse model is non-local. However, as we shall see in the next section, all the non-locality is relegated to inferring the pattern correlations, *ρ* and the two models are infact equivalent in the mean field limit.

#### 6.1 (Almost) equivalence of Gutfreund and Pseudo-Inverse model

In this section we show that the Gutfreund and pseudo-inverse model are (almost) equivalent when the stored patterns are hierarchically organized. We start by simplifying the overlap matrix in the mean field limit.

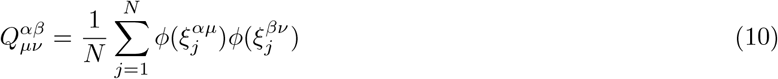

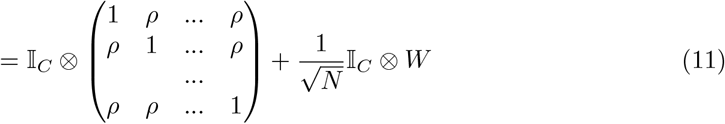

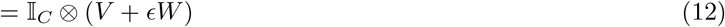

Here 𝕀_*C*_ is a *p*_*C*_ × *p*_*C*_ identity matrix, W is some *p*_*I*_ × *p*_*I*_ symmetric matrix and 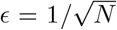. So, *Q*^−1^ = 𝕀 ⊗ (*V* + ϵ*W*) ^−1^. which can be calculated by Taylor expansion to first order in to give,

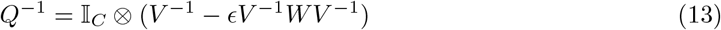

In the limit that *N, p* → ∞, the expression for *V* ^−1^ simplifies to give,

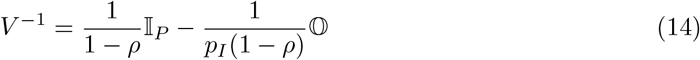

Here 𝕀_*P*_ is a *p*_*I*_ × *p*_*I*_ identity matrix, 𝕆 is a *p*_*I*_ × *p*_*I*_ matrix of all ones. Substituting *V* ^−1^ in the second term gives,

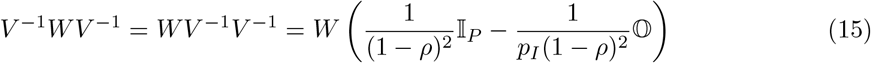

Let *W* 𝕆 = *p*_*I*_*k*𝕆 where *p*_*I*_*k* is some constant representing sum of row(or columns) of W matrix. So, the inverse of the overlap matrix can written as,

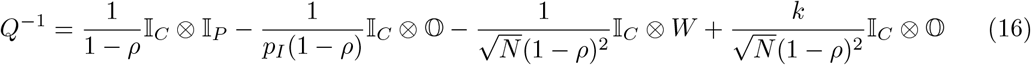

Substituting in the weight matrix,

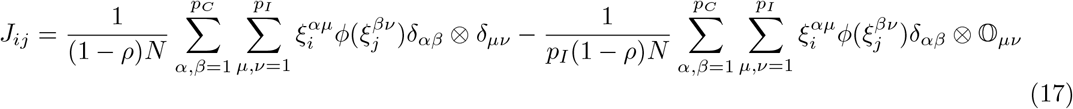

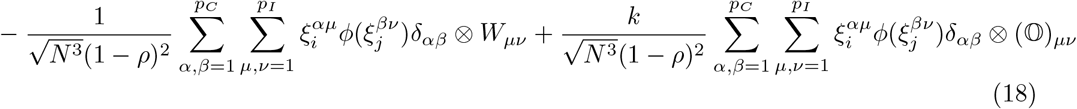

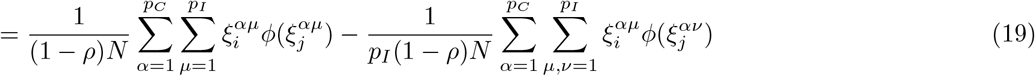

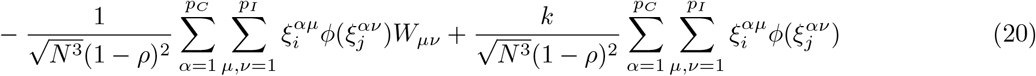

Let us set,

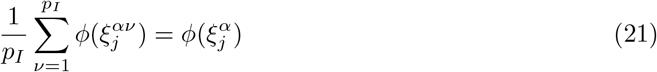

The weight matrix becomes,

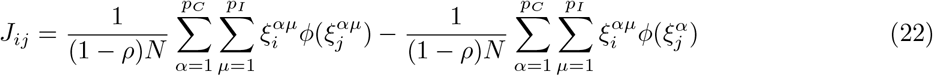

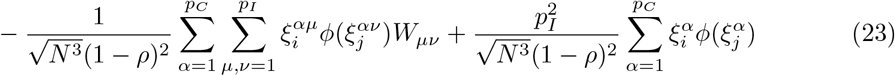

We notice that except for the third term the above expression is identical to Gutfreund model.

#### 6.2 Output correlation for different functional nonlinearities

The correlation between two random variables *ξ*_1_ and *ξ*_2_ is *ρ*. Then the correlation between *f* (*ϕ* (*ξ*_1_))) and *f* (*ϕ* (*ξ*_2_))) is given by the following expression:

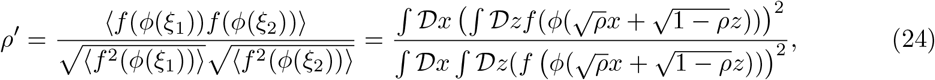

where 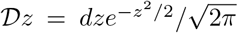. We want to find the relationship between the input and output correlation when *ρ* is small. Let *f* (*ϕ* (.)) = *g*(.). Then by Taylor expansion,

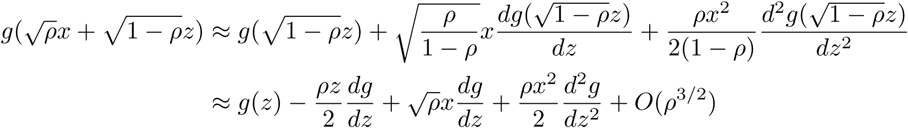

But for neurobiological reasons we want,

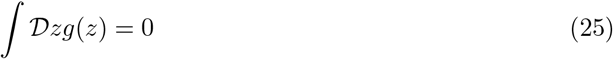

Consider the third term, we have,

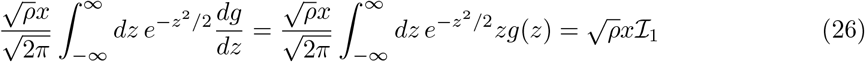

Where we have defined,

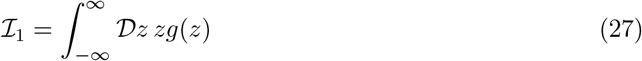

The denominator of the correlation for small values of *ρ* is given by:

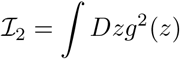

The output correlation integral becomes:

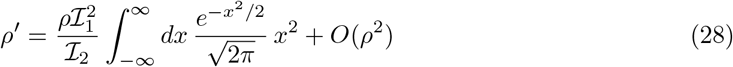

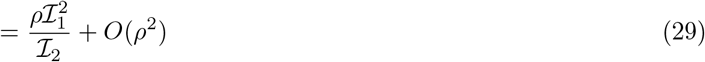

### 7 Mean Field Theory for hierarchically correlated patterns

#### 7.1 Gutfreund Model

We consider a neural network with *p*_*C*_ classes and *p*_*I*_ individual patterns per class. In the mean field limit, *p*_*C*_, *p*_*I*_ → ∞. The correlation structure is given by:

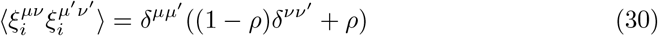

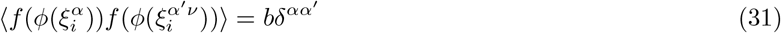

Here *µ* = 1, 2, …, *p*_*C*_ and *ν* = 1, 2, …, *p*_*I*_. The weight matrix is given by:

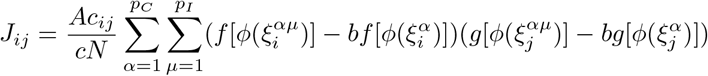

The input to neuron *i* is given by:

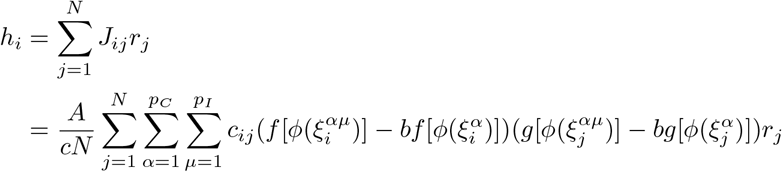

Let us assume the first pattern of the first class is being retrieved, *ξ*^11^. So, we separate the *α* = 1, *µ* = 1 term from other terms which we the call ‘noise term’

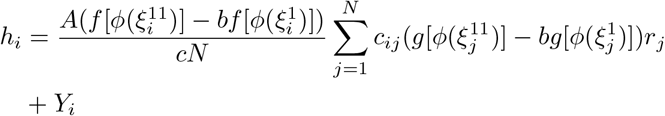

Here, the noise term *Y*_*i*_ is given by:

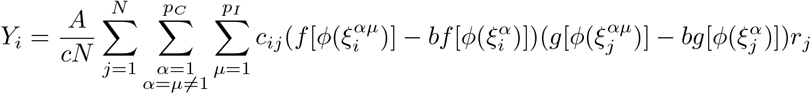

In the large *cN* limit, by the law of large numbers, the signal term of *h*_*i*_ simplifies to give:

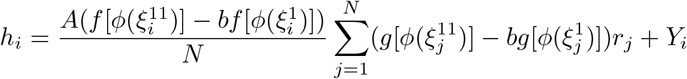

Let us define the order parameter q defined as following:

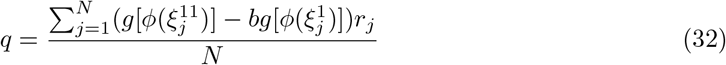

Using these definition, the expression for *h*_*i*_ becomes:

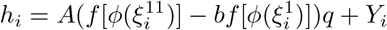

We now compute the statistics of the noise term. But first we write it in simplified notation:

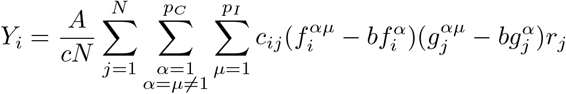

⟨*Y*_*i*_ ⟩= 0. Squaring *Y*_*i*_ gives,

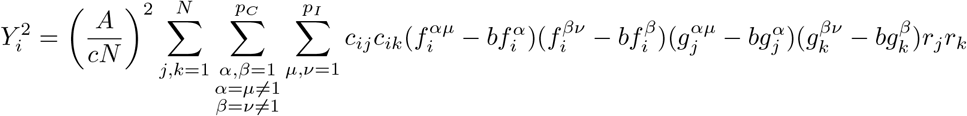

Taking the average with respect to *ξ*_*i*_,

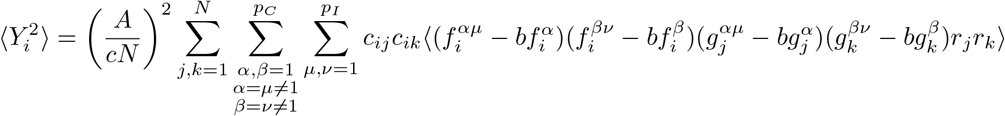

The cross terms corresponding to different neurons cancel out to give,

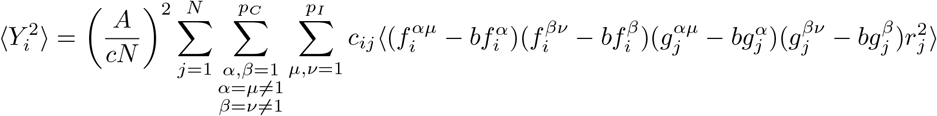

We now use law of large numbers to get,

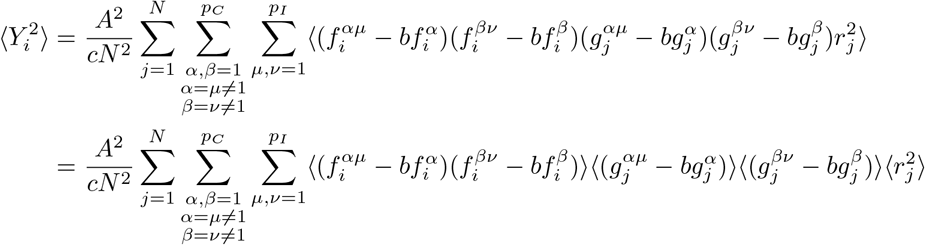

The second moment of 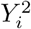 is given by:

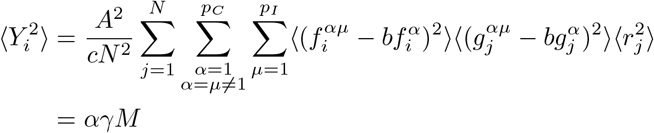

Here,

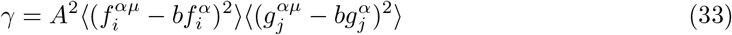

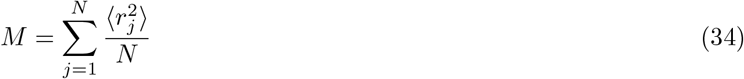

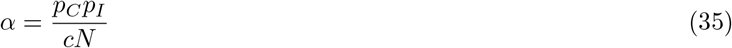

We now compute the order parameters self-consistently. The two equations are:

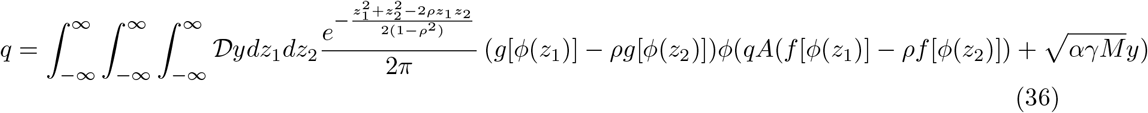

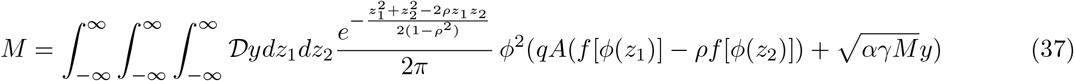

For solving these coupled integral equations, we consider the following system of differential equations:

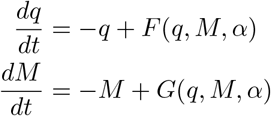

Here F and G correspond the right hand side of equation q and M equation respectively. The initial condition of the differential equation are *q*(*t* = 0) = *M* (*t* = 0) = 1.

The overlap m, which corresponds to the correlation between *g*(*ϕ* (*ξ*)) and the firing rates r, is given by

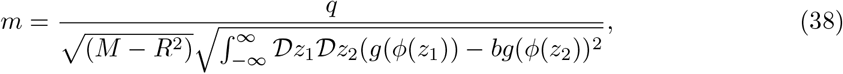

where R is the mean firing rate in the attractor state given by

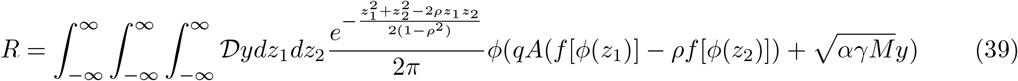

We plot m as function of *α* and mark the point of phase transition to get the maximum capacity.

#### 7.2 Pseudo-Inverse Model

As we saw in section 1.1, the weight matrix when the patterns are heirarchically organized is given by:

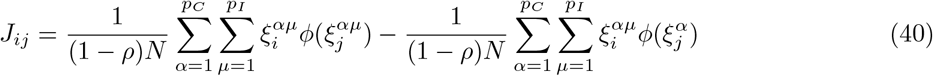

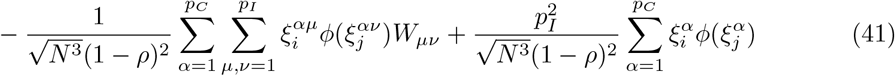

The input to neuron *i* is given by:

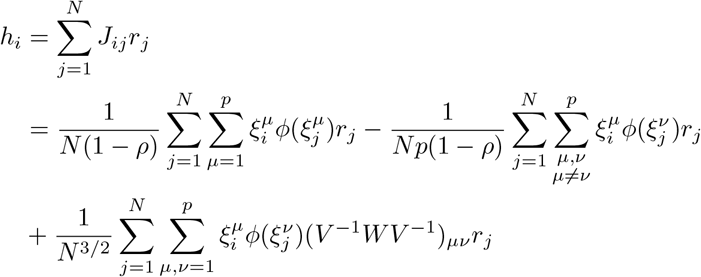

Let us assume the first pattern is being retrieved. So, we separate the *µ* = 1 from other terms to get:

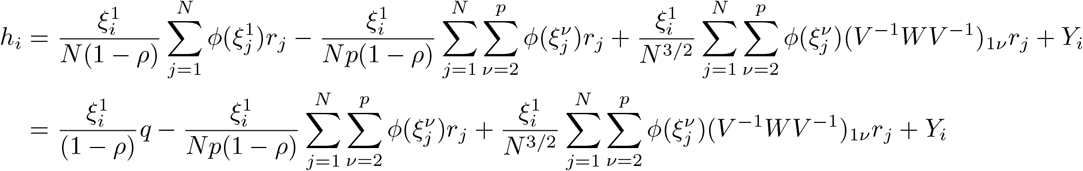

Where q is the order parameter as defined below:

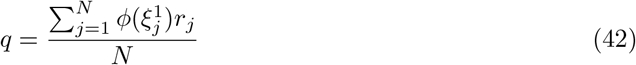

Here, the noise term *Y*_*i*_ is given by:

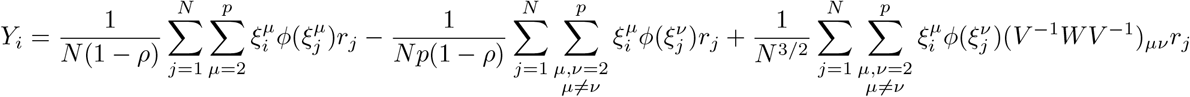

The next goal is to calculate the variance of *Y*_*i*_. Since, 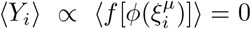, all we need to do is to calculate the second moment of *Y*_*i*_. To do that we first calculate the square of *Y*_*i*_,

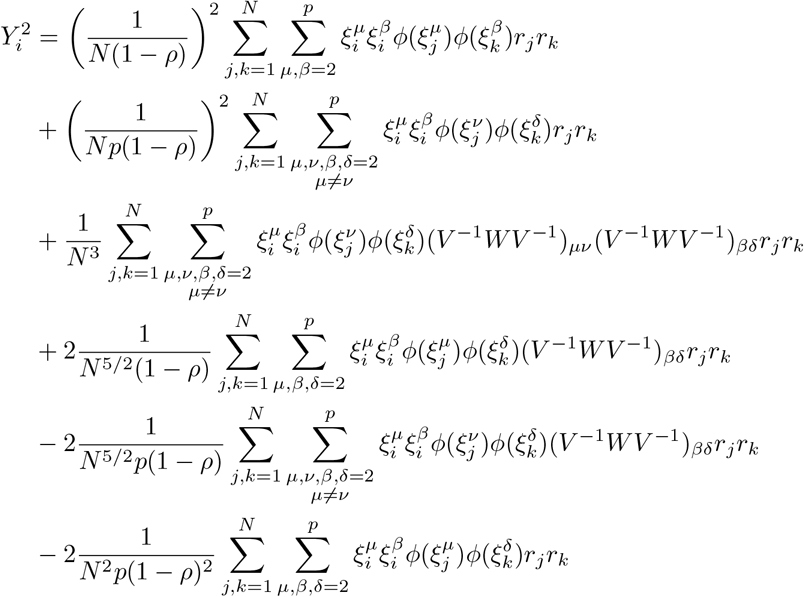

Taking the expectation value of *Y*_*i*_, using law of large numbers and cancelling out cross terms corresponding to different neurons gives us:

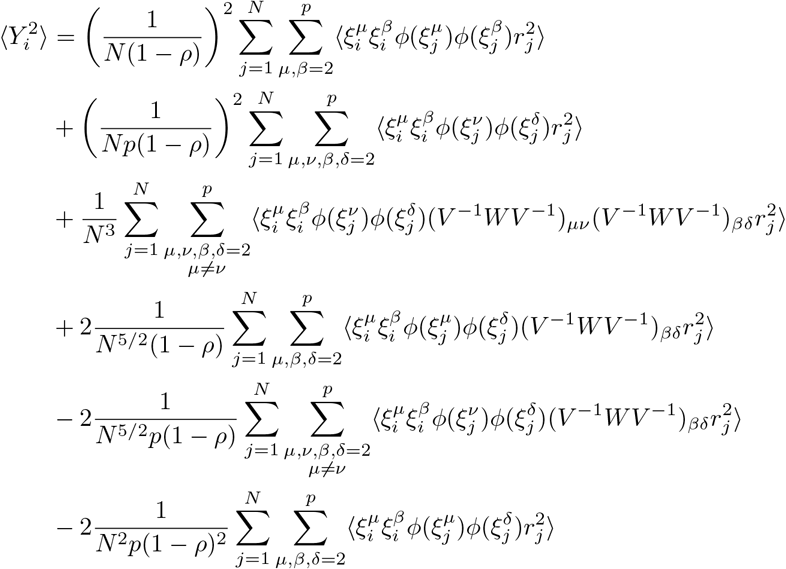

Since, we have 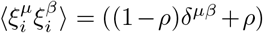, then 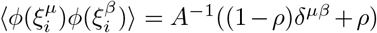. The expression for second moment of *Y*_*i*_ simplifies to:

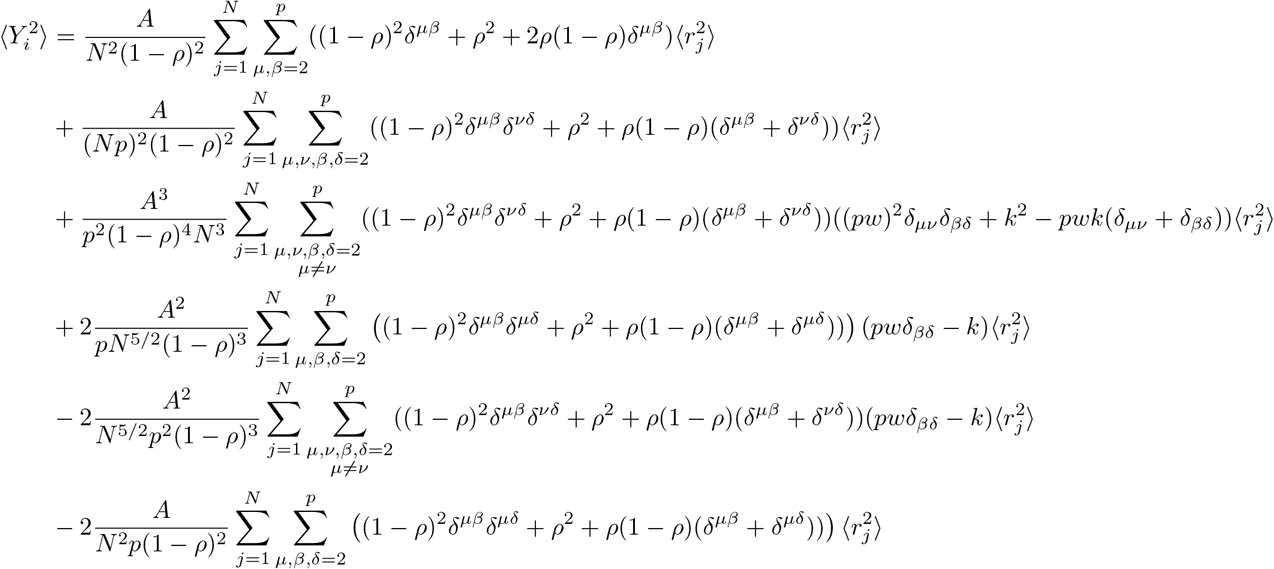

Let us define 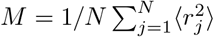. Substituting it the above expression and simplifying we get,

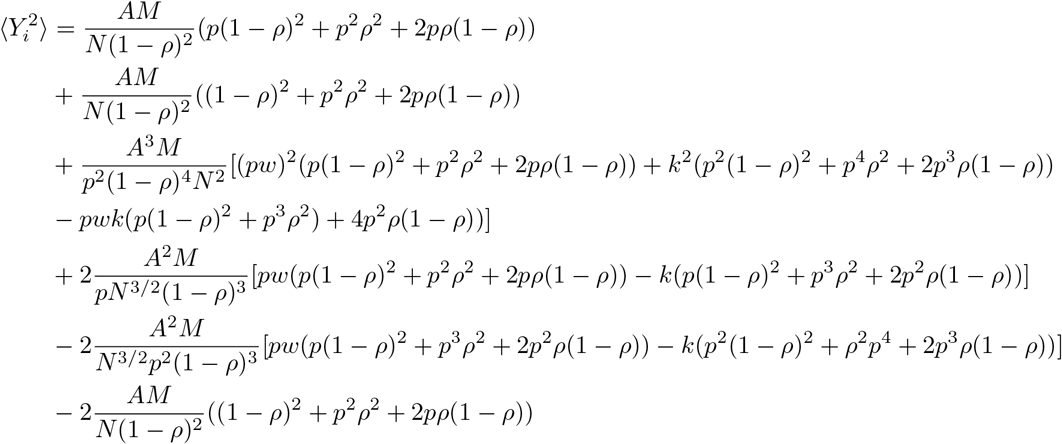

Combining the terms we get,

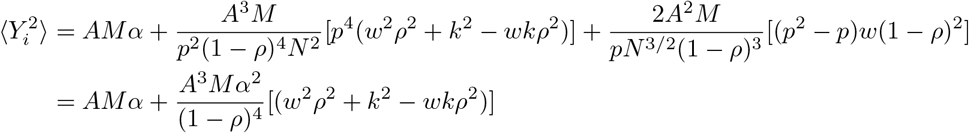

### 8 Excitatory-Inhibitory Rate network

We consider a network consisting of *N*_*E*_ excitatory neurons and *N*_*I*_ inhibitory neurons. The dynamics is given by the following system of differential equations:

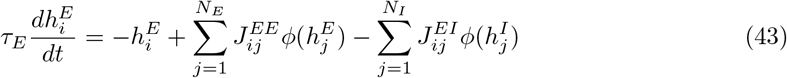

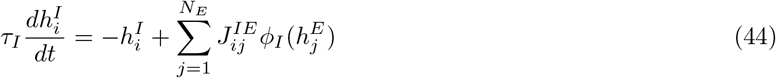

The weight matrix is given by:

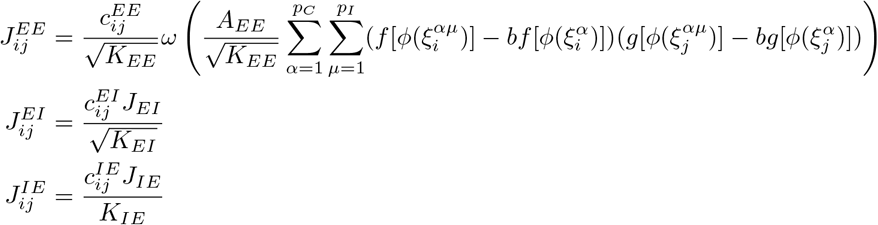

We now make the following assumptions:

1. Inhibition is much faster than excitation (*τ*_*I*_ ≪ *τ*_*E*_)
2. Inhibitory transfer function is linear. *ϕ*_*I*_ (*h*) = *gh*

With these assumptions, we just have one equation given by:

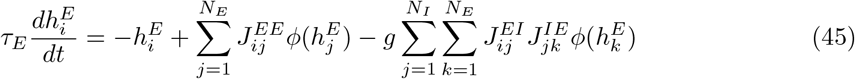

Taylor expanding 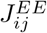, we get

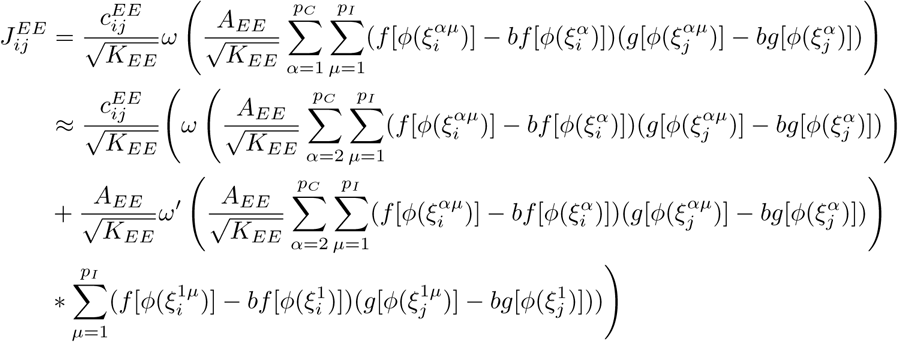

Substituting into 3 we get and averaging over *α* ≠ 1 we get:

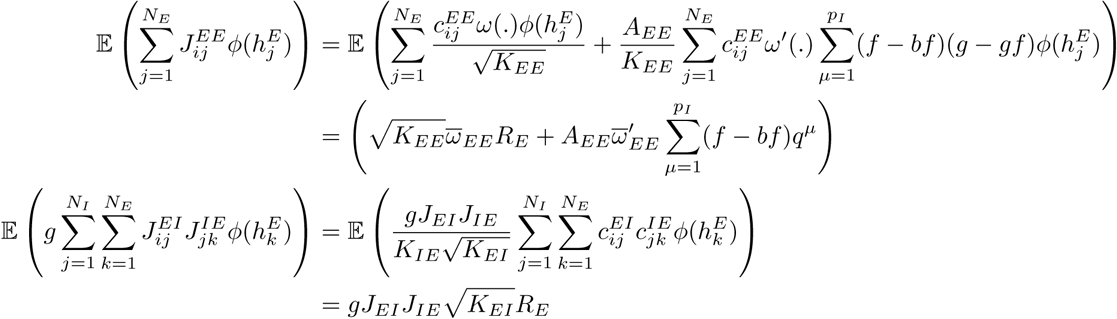

Where,

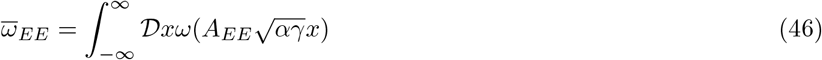

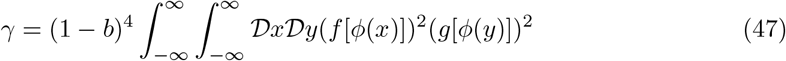

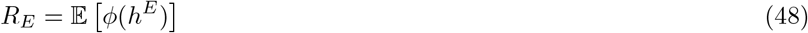

To get a balance between excitation and inhibition we impose the condition,

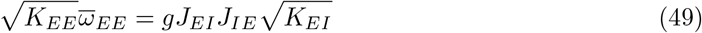

### 9 Excitatory-Inhibitory Spiking network

The EI spiking network is described by the following set of differential equations:

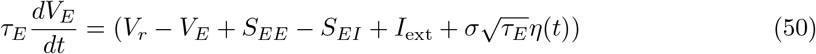

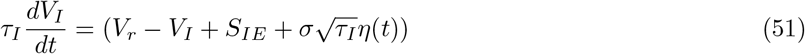

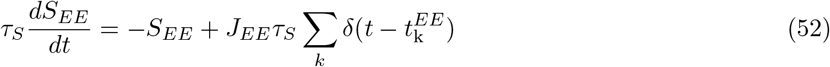

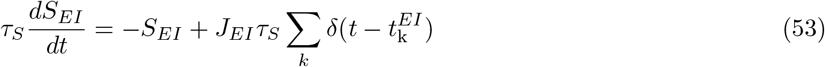

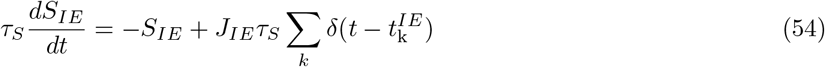

To go from EI rate network to EI spiking network we take the following steps:

- We want the mean firing rate to be the same for EI rate network and the EI spiking network. To do that, we fit the rate transfer function to LIF transfer function. There are two transfer function in the rate case *ϕ*_*I*_ (*x*) = *g*_*I*_*x* and *ϕ*_*E*_(*x*) which was of sigmoidal form. It turns out that given *V*_reset_, *V*_thrd_,*τ*_rp_ and gaussian noise of standard deviation *σ*, the firing rate of a particular spiking neuron is given by:

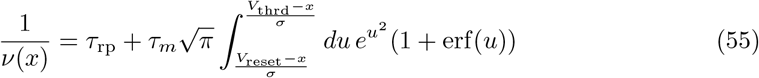

So, we need to figure out the parameters of the spiking neurons such that the mean firing rates are the same in rate and spiking case. To do that, we use L-BFGS-B algorithm.

### 10 Parameter Tables

#### Plasticity Rule Parameters

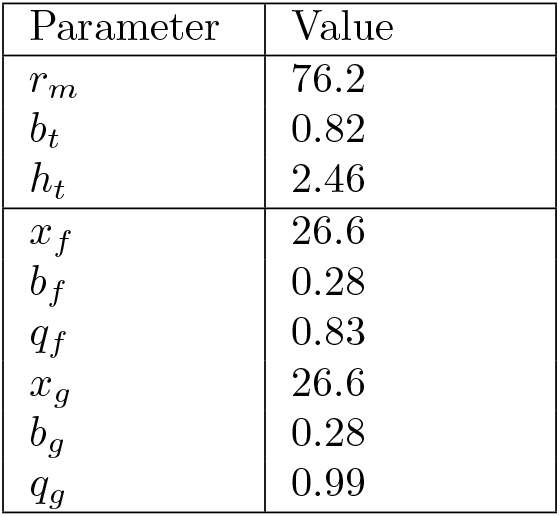

#### One population rate network

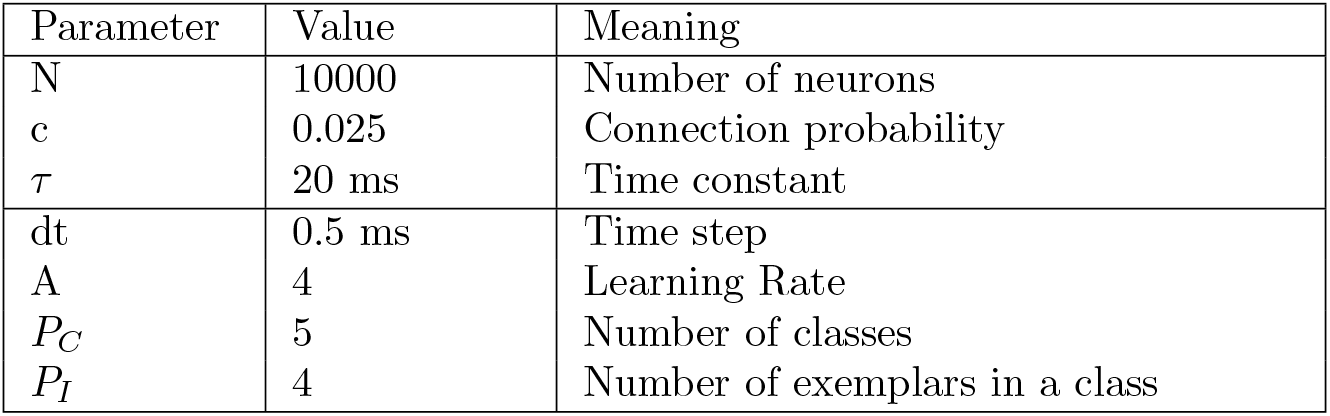

#### Two population rate network

| Parameter | Value | Meaning |
| --- | --- | --- |
| $N_E$ | 10000 | Number of excitatory neurons |
| $N_I$ | 2500 | Number of inhibitory neurons |
| $c_{EE}$ | 0.05 | E to E connection probability |
| $c_{EI}$ | 0.02 | I to E connection probability |
| $c_{IE}$ | 0.02 | E to I connection probability |
| $\tau_E$ | 20 ms | Excitatory time constant |
| $\tau_I$ | 5 ms | Inhibitory time constant |
| dt | 0.5 ms | Time step |
| $A_{EE}$ | 6 | Learning Rate |
| $P_C$ | 5 | Number of classes |
| $P_I$ | 4 | Number of exemplars in a class |
| $J_{EI}$ | 0.0250 | I to E synaptic weight |
| $J_{IE}$ | 0.0673 | E to I synaptic weight |

#### Two population spiking network

| Parameter | Value | Meaning |
| --- | --- | --- |
| $N_E$ | 10000 | Number of excitatory neurons |
| $N_I$ | 2500 | Number of inhibitory neurons |
| $c_{EE}$ | 0.05 | E to E connection probability |
| $c_{EI}$ | 0.02 | I to E connection probability |
| $c_{IE}$ | 0.02 | E to I connection probability |
| $\tau_E$ | 20 ms | Excitatory time constant |
| $\tau_I$ | 5 ms | Inhibitory time constant |
| $\tau_S$ | 20 ms | Synaptic time constant |
| dt | 0.5 ms | Time step |
| $A_{EE}$ | 6 | Learning Rate |
| $P_C$ | 5 | Number of classes |
| $P_I$ | 4 | Number of exemplars in a class |
| $J_{EI}$ | 0.0250 | I to E synaptic weight |
| $J_{IE}$ | 0.0673 | E to I synaptic weight |
| $V_r$ | -65 mV | Resting/Reset Voltage |
| $V_{th}$ | -50 mV | Threshold Voltage |
| $V_{min}$ | -75mV | Minimum Voltage |

